# Mechanical compression induces neuronal apoptosis, reduces synaptic activity, and promotes glial neuroinflammation in mice and humans

**DOI:** 10.1101/2025.05.22.655651

**Authors:** Maksym Zarodniuk, Anna Wenninger, Julian Najera, Jihaeng Lee, Jack Markillie, Cameron MacKenzie, Jenny Bergqvist-Patzke, Bianca Batista, Megna Panchbhavi, R’nld Rumbach, Alice Burchett, Meenal Datta, Christopher Patzke

**Affiliations:** Department of Aerospace and Mechanical Engineering; Department of Biological Sciences; Department of Chemistry and Biochemistry; Department of Chemical and Biomolecular Engineering; Department of Applied and Computational Mathematics and Statistics, University of Notre Dame

## Abstract

Mass effect, characterized by the compression and deformation of neural tissue from space-occupying lesions, can lead to debilitating neurological symptoms and poses a significant clinical challenge. In the primary brain tumor glioblastoma (GBM), we have shown previously that compressive solid stress originating from the growing tumor reduces cerebral blood flow, leads to neuronal loss, increased functional impairment, and poor clinical outcomes. However, the direct effects of compression on neurons and the underlying biophysical mechanisms are poorly understood. Here, using multi-scale compression systems and physiologically relevant *in vitro* and *in vivo* models, we find that mechanical compression induces neuronal apoptosis and synapse loss, leading to disrupted neural network activity. This is accompanied by increased HIF-1 signaling and upregulation of downstream stress-adaptive genes in neurons. We further show that compression triggers AP-1–driven gene expression in glial cells, promoting a neuroinflammatory response. Together, these findings reveal that solid stress directly contributes to neuronal dysfunction and inflammation caused by GBM by activating distinct pathways that can be targeted in future studies for neuroprotection.

**SIGNIFICANCE STATEMENT:** Glioblastoma (GBM), the deadliest primary brain tumor in adults, exerts physical forces on surrounding brain tissue as it grows, leading to neuronal damage. However, the molecular mechanisms underlying this process are not well understood. In the present study, by applying multiple model systems, we show that mechanical compression triggers neuronal apoptosis, disrupts synaptic communication between neurons, and reduces neural network activity. We also find that compression activates inflammatory pathways in both neurons and glia, further contributing to neuronal damage. These findings reveal how compression exerted by space-occupying lesions may contribute to patients’ cognitive and motor impairments and suggest new directions for treatment. This work lays the groundwork for therapies that protect neurons from mechanical injury, with relevance not only to GBM but also other neurological diseases that present with mass effect.

## INTRODUCTION

Several central nervous system pathologies such as traumatic brain injury (TBI), multiple sclerosis (MS), benign and malignant brain tumors are known to present with mass effect, where a focal lesion or contusion compresses and deforms surrounding neural tissue due to the space occupied by increased cell numbers (arriving via circulation and/or proliferating locally) and/or extracellular matrix (ECM) deposition within a confined area. Such deformations, including ventricular compression and displacement of the brain midline, are immediately visible on magnetic resonance imaging (MRI). Clinicians frequently observe both fluid (i.e., edema, resulting from disrupted or leaky vasculature and/or cerebrospinal fluid accumulation) and solid components of these lesions and evaluate these criteria during diagnosis (1–3). Intracranial mass effect (as opposed to compression of the spinal cord or peripheral nervous system) is particularly unique and insidious, as the rigid skull confines and compounds these mechanical forces (4). As a result, patients often present with debilitating neurological symptoms including sensory, motor, cognitive, behavioral, and emotional dysfunction (5, 6).

In the case of glioblastoma (GBM), compressive solid stresses are generated by solid and elastic components of the tumor microenvironment, namely the cells and ECM (7–9). Additionally, peritumoral edema promotes increased intracranial pressure and further deforms brain tissue (10). Similar space-occupying lesions, presenting with an overlapping but distinct set of symptoms have been observed in other neurological disorders such as tumefactive MS (11, 12) and severe TBI (13–15). Both edema and solid stress contribute to brain deformation and have been measured/estimated and investigated as a prognostic/predictive biomarker in brain tumor and TBI (2, 4, 16, 17). In GBM, the extent of peritumoral edema is an independent prognostic factor; however, its direct effect on neurological performance is unclear (4, 18). In contrast, solid stress has been linked to increased functional impairment, reduced cerebral blood flow (CBF), and poorer clinical outcomes in GBM patients (4, 19). Our prior preclinical investigations have shown that chronic brain compression using mechanical compression techniques *in vivo* lead to similar effects, such as reduced vessel perfusion, neuronal loss, and neurological dysfunction, even in the absence of a tumor (4).

Growing brain masses, such as tumors, cause tissue compaction that generates both compressive and tensile forces. These mechanical stresses can compress, shear, and stretch axons, triggering a series of cellular processes that ultimately contribute to neuronal dysfunction and death (20). In TBI, acute mechanical damage has been shown to trigger long-term dysregulation of intracellular calcium metabolism (21, 22), excitotoxicity (23), Wallerian degeneration (24), and neuroinflammation (25, 26) in injured neurons. While these mechanisms are well-documented in the context of acute injury, the effects of chronic compressive forces, such as those produced by a growing tumor, remain less understood.

In this study, we utilize compression systems to examine the direct effects of mechanical compression on neurons. We show that chronic compression of iPSC-derived human neurons *in vitro* leads to neuronal and synaptic loss, as well as impaired neural network activity. Additionally, we find that compression activates reactive gene expression programs in astrocytes and microglia, promoting neuroinflammatory damage through secretion of inflammatory cytokines. Our findings provide new insights into how solid stress contributes to neuroinflammation and neurodegeneration in GBM and other neurological diseases and inform future strategies to reduce these deficits in patients suffering from mass effect.

## METHODS

### Human induced neurons (iN cells) and lentivirus generation

Induced pluripotent stem cells (iPSCs) from three different healthy wild-type donors were used for experiments: WB02 (Asian male, 25-29 years) referred to iPSCs #1 UKKi027-A (African female, 25-29 years) referred to iPSCs #2, and ATCC-DYS0100 (ACS-1019; from neonate male) referred to iPSCs #3. Induced neurons were produced following our previously published standard protocol (27, 28). Briefly, iPS cells were grown on 6-well dishes coated with Matrigel (Corning) in Stem Flex (Thermo Fisher) supplemented with rock inhibitor (thiazovivin, Bio Vision,1:5000). Media was changed daily. When iPSCs were forming colonies and becoming confluent, 1 mL accutase (Innovative Cell Technologies) was used to dissociate cells from the plate. Cells were put in the incubator for around 5 minutes. Cells were collected with 1 mL of media in a 15 mL conical tube and centrifuged for 5min at 300 x g. The supernatant was removed, and the cell pellet was resuspended in 1 mL Stem Flex. 10uL of the resuspension was used to count cells using the BioRad automated cell counter. For each cell line, 120k cells were plated on 6-well 0.4 µm PET translucent transwell inserts (cellQUART) that had been coated in Matrigel for 24 hours. Cells used for live imaging were lentivirally transduced with 110 uL FUW-rtTA, 110 uL TetO-Ngn2, 110 uL FUW-GCamp8m (modified after pGP-CMV-jGCaMP8m, which was a gift from GENIE Project (Addgene plasmid # 162372 ; http://n2t.net/addgene:162372 ; RRID:Addgene_162372) (29), and 110 uL FUW-GFP. Cells used for RNA sequencing were lentivirally transduced with 110 uL rtTA and 110 uL Ngn2. The mastermixes included counted cells, lentivirus, thiazovivin (1:5000), and Stem Flex. 1.5mL of mastermix was plated onto each transwell. On day 0, the culture medium was replaced with DMEM/F12 (Thermo Fisher) supplemented with 1X N2, 1X NEAA (Thermo Fisher) containing human BDNF (10 ng/ml, PeproTech), human NT-3 (10 ng/ml, PeproTech), and mouse Laminin-1 (0.2 µg/ml, Thermo Fisher). Doxycycline (2 µg/ml, Sigma) was added on day 0 to induce TetO gene expression and retained in the medium until the end of the experiment. On day 1, a 48h puromycin selection (1 µg/ml, InvivoGen) period was started. On day 3, mouse glia cells were added in neurobasal-A medium (Thermo Fisher) supplemented with B27/Glutamax (Thermo Fisher) containing Ara-C (2 µM, Sigma). After day 3, 50% of the medium in each well was exchanged every 2 days. FBS (5%, Corning) was added to the culture medium on day 5 to support astrocyte viability, and human neurons were assayed after 2 months (Figure 1A-B).

**Figure 1.**
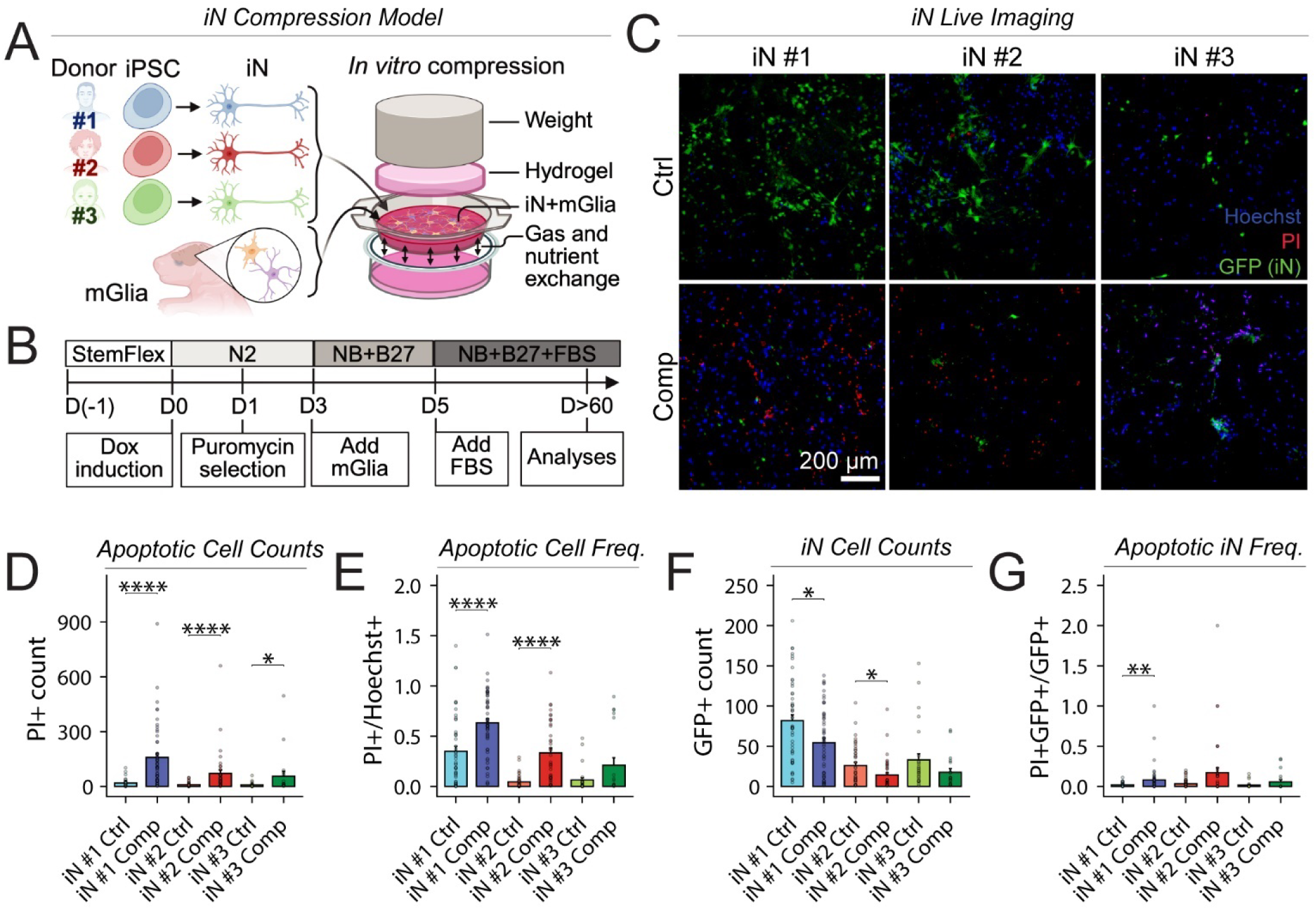
Compressive solid stress induces neuronal cell death in human Ngn2-induced neurons. (**A**) Schematic of the *in vitro* compression system used to apply 0.14 kPa of compressive stress on human induced excitatory neurons (iNs), mimicking solid stress exerted by a GBM tumor. iNs are derived from human induced pluripotent stem cells (iPSCs) from diverse donors and are co-cultured with murine glia (mGlia) to facilitate neuron maturation and synapse formation. (**B**) Diagram showing the protocol used for iN induction from iPSCs. (**C**) Representative images of iN cultures stained with propidium iodide (PI) and Hoechst 33342 after 24-hour compression. GFP expression in iNs is largely restricted to the nucleus. (**D**) PI+ (non-viable) cell numbers are quantified as the number of PI-positive particles per field of view. (**E**) PI+ cell frequency is defined as the number of PI+ cells divided by the total cell number determined by Hoechst 33342 staining. (**F**) Number of GFP+ neurons is quantified as the number of GFP+ particles. (**G**) PI+ (apoptotic) neuron frequency is quantified as the number of PI and GFP double-positive foci normalized to GFP+ neuron number. Mann–Whitney *U* test with Holm– Bonferroni adjustment for multiple comparisons using N = 4 biological replicates for iPSCs #3, n=7 biological replicates in iPSCs #2 and iPSCs #1 neurons per group; the data are based on 7 independent experiments; Error bars: mean ± SEM. Due to puncta-based quantification, some frequencies may exceed 1. Statistical significance is indicated as follows: ns = P > 0.05; * = P ≤ 0.05; ** = P ≤ 0.01; *** = P ≤ 0.001; **** = P ≤ 0.0001.

Lentiviruses were produced in HEK293T cells (ATCC, VA) by co-transfection with three helper plasmids (pRSV-REV, pMDLg/pRRE and vesicular stomatitis virus G protein expression vector) with 12 µg of lentiviral vector DNA and 6 µg of each of the helper plasmid DNA per 75 cm^2^ culture area) using calcium phosphate. 12 mL of lentivirus-containing supernatant was harvested 48 h after transfection, aliquoted, and frozen at -80°C (30).

### Astrocyte and microglia cell culture

Immortalized human astrocytes (IHA, Creative Bioarray) were seeded at 200,000 cells per insert in 1 mL of DMEM supplemented with 5% FBS (Thermo Fisher), and 1X Pen/Strep (Corning) (31). Human C20 microglia (32–34) were seeded at 20k cells in 0.4 μm pore size transwell cell culture insert (CellQART, 4.5 cm²) in 1 mL of DMEM/F12 media supplemented with 1X N2 (Thermo Fisher), 1% FBS, and 1X Pen/Strep. Once adherent, the cells were compressed for a defined period of time. For live cell morphology analysis, the cells were washed twice with 1x PBS, stained with 0.8 µg/ml Calcein AM (BioLegend) and 12 µg/ml PI (MP Biomedicals) diluted in 1 ml of 1x PBS for 30 minutes at 37°C, and washed twice again with 1x PBS. Images were taken at 20x magnification using a fluorescent microscope (Leica DMi8) and analyzed in ImageJ Software. Microglia circularity was analyzed by converting images to binary, de-speckling, thresholding to obtain masks of the soma region, and measuring circularity of individual cells. Soma lengths and widths were measured manually using the ImageJ measuring tool.

### *In vitro* compression

To apply a defined amount of compressive stress, we used a previously described *in vitro* compression device (35). Briefly, a 1% agarose cushion was placed on top of the cells in the transwell insert, and a 3D-printed PLA weight was placed on top to apply 0.14 kPa of compressive stress for 24 hours. This level of solid stress is meant to simulate the solid stress that we previously measured in murine GBM models (4). Control cells were covered with an agarose cushion only (0 kPa).

### Live/Dead staining

Cell viability was assessed using Hoechst 33342 (Invitrogen) and Propidium Iodide (PI). Hoechst 33342 is a cell-permeant nuclear dye and was used for data normalization. PI is a membrane impermeant dye and stains apoptotic and/or necrotic cells whose cell membrane is disrupted. To allow for discrimination of neurons vs glia, we overexpressed GFP in neurons. After weights and cushions were carefully removed together with any remaining media, 0.5 mL of working solution containing 3 µg/mL PI and 1:1000 Hoechst 33342 diluted in the imaging buffer (4mM CaCl2 and 8mM KCl in 129 mM NaCl, 25 mM HEPES, 30mM glucose, 1 mM MgCl2, 10 µM Glycine) were added to the transwell insert, and 1 mL was added underneath the insert into the cell culture well. The cells were allowed to incubate for 10 min at 37°C, 5% CO^2^ before imaging.

### Calcium Imaging

Following compression or control treatment, the cells were used for calcium imaging. The weight was carefully removed from the compressed neurons. Existing media was aspirated from inside and outside of the transwell. Calcium imaging was performed as previously described (36, 37) and modified for agarose-covered transwell condition. Briefly, after lentivirally mediated expression of GCaMP8m for 2 months, 1mL of calcium imaging buffer (4mM CaCl2 and 8mM KCl in 129 mM NaCl, 25 mM HEPES, 30mM glucose, 1 mM MgCl2, 10 µM Glycine) was pipetted outside the transwell, and 200uL on to the agarose cushion in the transwell. Neurons were placed in the incubator for 3-5 minutes and then imaged in the transwells with an inverted epi-fluorescence microscope (Nikon EclipseTS2R) with a 488 nm filter at room temperature. GCaMP8m fluorescence was recorded for 2 min at a frame rate of 50ms using NIS Elements 5.30.05. The ROI was set to 1192x1192 Mono 16-bit, the objective was 10x. For each condition, 7 fields of view were imaged for each biological batch and condition. All images were acquired using the same exposure time. Time-lapse imaging was performed in areas containing confluent neuron populations with few overlapping cell somata. This procedure was used for compressed and uncompressed samples.

Calcium imaging files were analyzed using MATLAB through custom-made code. Neurons were selected and then each were analyzed to generate statistics summary: frequency and mean/standard deviation of amplitudes for each neuron and synchronous spikes (dependent on the sigma threshold we set, 1.5). For each replicate, cell line, and compressed/uncompressed, the data was combined and graphed using GraphPad Prism: Amplitude, Frequency, Synchronous Amplitude, and Synchronous Frequency.

### Confocal Imaging and Immunocytochemistry (ICC)

After 60 days in culture, human induced neurons were collected for ICC and confocal imaging. The agarose cushions and weights were carefully removed from the transwells. The neurons were fixed in 4% paraformaldehyde and 4% sucrose in PBS for 20 min at room temperature, washed three times with PBS. Cells were blocked in PBS containing 5% goat serum and 0.2% Triton X-100 for 1 h at room temperature and stored in the blocking buffer for up to 2 weeks. Primary antibodies were applied overnight at 4°C, cells were washed in PBS three times, and fluorescent-labelled secondary antibodies (Alexa, 1:1000) were applied for 2 h at room temperature in the dark. The following antibodies were used in immunocytochemistry experiments: MAP2 (CPCA-MAP2, EnCor. 1:5000), synapsin (7739; 1:2000). The transwells were carefully washed 3x with PBS and 1x with ultrapure water. The membranes of the transwells were carefully removed and cut into a square. The membrane containing neurons was mounted onto a coverside with DAPI fluoromount-G (Southern Biotech) media and a square coverslip. Slides were left to dry for 24 hours.

Confocal imaging was performed using a Nikon Ti2 eclipse A1RSi confocal microscope using NIS elements version 5.30.06. 64-bit. 1024 x 1024 images were taken with 60x objective with 0.8µm step size and 7-10 steps per image, exported as .nd2 files and uploaded to NIS elements version 5.42.04 64-bit. Each image was analyzed by making a new document from the current view with Maximum Intensity Projection (MaxIP). The GFP threshold was defined as 1000 - 4095, Clean 0.11um, Separate 0.11um. Individual regions of interest (ROIs) were drawn around dendrites (10 ROIs per image). Binary ROI data was collected with Automated Measurement Results and exported to Microsoft Excel. To measure the length of the dendrite in µm, we used the polyline tool in Annotations and Measurements. Puncta density was calculated by dividing the Number of Objects by the length of dendrite. Puncta area was calculated by dividing the Binary Area by the Number of Objects. Puncta intensity was calculated by dividing the Sum Intensity by the Number of Objects.

Confocal immunohistochemistry (IHC) images were analyzed using ImageJ (version 1.54g). Background signal was removed from all channels using the “Remove Background” function with a radius of 100. To eliminate regions of non-specific vascular staining, pixel intensities were capped at a predefined threshold that was applied consistently across all images and experimental groups. The percentage area of GFAP+ staining was calculated as the GFAP+ area divided by the total DAPI+ area. NeuN+ cell frequency was quantified by dividing the number of NeuN+DAPI+ particles by the total number of DAPI+ particles. To quantify nuclear HIF1A mean fluorescence intensity (MFI), masks were generated for DAPI+ nuclei that were also NeuN+, corresponding to neuronal nuclei. MFI was measured for each particle, and the average was calculated for each biological replicate (i.e., each animal). The frequency of HIF1A+ neurons was determined by dividing the number of HIF1A+NeuN+DAPI+ particles by the total number of DAPI+ particles.

### Hypoxia evaluation

Prior to compression, cells were cultured in transwell inserts and incubated overnight with 100 nM HypoxiTRAK™ (Novus Biologicals) to allow dye uptake. The following day, cells were subjected to mechanical compression by placing an agarose gel and a weight into the insert for 24 hours. After the weight was removed, the cells were gently washed with phosphate-buffered saline (PBS) and detached using Trypsin-EDTA (0.25%). Following detachment, cells were washed with PBS. Zombie Aqua™ viability dye (BioLegend) was diluted 1:400 in PBS, and cells were resuspended in 50 µL of the diluted dye. Samples were incubated at 4°C in the dark for 15–30 minutes, then washed once with 1% bovine serum albumin (BSA, VWR) in PBS. Cells were subsequently fixed in 100 µL of 4% paraformaldehyde (PFA) for 10 minutes at room temperature, washed again, and immediately analyzed by flow cytometry (Cytek Aurora).

### PCR

Following compression, cells were lysed in 350 μl of TRI-reagent (Zymo Research, R2050-1-200) and the RNA was purified using an RNA isolation kit (Zymo Research, R2051). Gene expression was analyzed using TaqMan primers for Fosl2 (Hs01050117_m1), Fosl1, (Hs00759776_s1), Tnf, (Hs00174128_m1), Gapdh (Hs02786624_g1), Fos (Hs04194186_s1), Col4a1 (Hs00266237_m1), Il1b (Hs00266237_m1), Fosb (Hs00171851_m1). The raw PCR data was analyzed using the qPCR Design and Analysis app (Thermo Fisher Scientific). Individual measurements that were clear outliers (e.g. due to pipetting errors) were removed.

### RNA sequencing

RNA-seq library preparation and sequencing was performed at the Notre Dame Genomics and Bioinformatics Core Facility. Quality of RNA samples was confirmed using High Sensitivity RNA ScreenTape (Agilent) before Illumina libraries were prepared from 200 ng of RNA using the NEBNext Ultra II Directional RNA Library Prep kit with Sample Purification Beads (E7765S) and using the NEBNext Poly(A) mRNA Magnetic Isolation Module (E7490S). Libraries were confirmed using the Fragment Analyzer and quantified by qPCR prior to pooling and sequencing on an Illumina NextSeq500 using 50+50 paired end reads. On average, 1 NextSeq P4 XLEAP flow cell generates 1.5-1.8 Billion clusters/reads, for an average of 125 Million raw reads per sample across both lanes.

### Analysis of iN RNA-seq data

To deconvolve neuronal and glial transcripts, sequencing reads were binned by mapping to human (GRCh38.p14) and mouse (GRCm39) reference genomes using BBMap (v. 39.08). Reads that mapped to both references were discarded. Binned reads were aligned to respective reference genomes and counted using STAR (v. 2.7.2). For PCA, gene counts were downsampled to the same library size and batch effects were regressed out using removeBatchEffects function in limma (v. 3.62.2) R package. The Bioconductor R package DESeq2 (v. 1.46.0) was used to normalize count data and perform differential expression analysis. Gene set enrichment and over-representation analysis was performed using clusterProfiler R package.

To infer the relative abundances of murine cell types in the mouse glial RNA-seq data, we applied non-negative least squares (NNLS) regression using the top 100 marker genes for each cell population from the Tabula Muris scRNA-seq dataset. First, mouse brain scRNA-seq data was normalized and clustered using SNN-based clustering in Seurat (v. 5.2.1). Marker genes for each cell population were identified through the Wilcoxon Rank Sum test. Pseudo-bulk aggregate expression profiles for 100 marker genes based on avg_log2FC were normalized, and relative abundances of the cell populations in the bulk RNA-seq data were inferred using NNLS, implemented via the lsei R package (v. 1.3-0). Motif enrichment analysis was performed using HOMER (v5.1) with the findMotifs module, scanning regions from 400 bp upstream to 100 bp downstream of the transcription start site (TSS). Motif lengths of 8 and 10 base pairs were used for the analysis.

### Analysis of Ivy GAP RNA-seq data

Anonymized BAM files were downloaded from the Allen Brain Atlas API and were counted using featureCounts (v2.0.1) and GRCh37.p5 reference genome. DESeq2 (v. 1.46.0) was used to normalize count data and perform differential expression analysis. Neuronal gene expression deconvolution was performed using BayesPrism (v. 2.2.2) and scRNA-seq reference (38). To pre-process scRNA-seq data, the count matrix was log-normalized, scaled, and clustered using the shared nearest neighbor (SNN)-based algorithm in FindClusters Seurat (5.1.0) function. The clusters were annotated based on marker genes provided in the parent publication (38). For gene expression deconvolution, genes located on the sex chromosomes were excluded from the reference to avoid sex-specific transcriptional states, and non-coding, ribosomal protein-coding as well as mitochondrial genes were removed to minimize batch effects. Inferred gene expression values were rounded to the nearest integer and normalized using the DESeq2 workflow.

### Analysis of Ivy GAP MRI data

Midline shift (MLS) was measured using a previously described method (39). MLS was defined as any deviation of the septum pellucidum from the midline. MLS was calculated by drawing a line between the anterior and posterior falx cerebri, then measuring the perpendicular distance to the point of maximal deviation from the septum pellucidum. Multivariate linear regression between MLS and tumor, edema volume, and several tumor shape descriptors revealed a positive association with edema volume. To separate the effects of fluid pressure (edema) from solid stress (e.g., mass effect from solid components of the tumor), we regressed MLS on edema volume and used the residuals as a proxy for solid stress. Patients with positive residuals—indicating more shift than expected from edema alone—were classified as “high” solid stress; those with zero or negative residuals were classified as “low. To study gene expression at the tumor margin in these two groups, we analyzed RNA-seq and ISH data from histologically annotated “leading edge” regions, where the malignant-to-normal cell ratio is approximately 1–2 per 100 (40).

### Mice

Animal experiments conducted in this study were in accordance with National Institute of Health Guidelines for the Care and Use of Laboratory Mice and approved by the Institutional Animal Care and Use Committee (IACUC) at the University of Notre Dame. Mouse glial cells were cultured from the forebrain of newborn wild-type CD1 mice. Briefly, newborn mouse forebrain homogenates were digested with papain and EDTA for 15 min, cells were dissociated by harsh trituration to avoid growing of neurons, and plated onto T75 flasks in DMEM supplemented with 10% FBS. Upon reaching confluence, glial cells were trypsinized and replated at lower density once to remove potential trace amounts of mouse neurons before the glia cell cultures were used for co-culture experiment with human neurons.

### *In vivo* compression model

To assess the impact of cranial compression on neuronal viability, cranial compression windows (cCWs) were surgically implanted in adult C57BL/6 mice, as previously described (4, 9). Compression was applied daily for three weeks by tightening the screw 67 degrees per day, resulting in an estimated volumetric displacement of approximately 1.3 mm³/day. At the study endpoint, mice were euthanized and the entire head was fixed in paraformaldehyde at 4°C for 48 hours. Brains were then dissected, rinsed in phosphate-buffered saline, and stored until further processing. Fixed tissues were processed using standard histological protocols and embedded in paraffin.

### Tissue Immunohistochemistry (IHC)

Paraffin-embedded tissue sections (10 µm) were deparaffinized in xylene and rehydrated through a graded ethanol series (100%, 95%, 70%, 50%, and 0%). Antigen retrieval was performed by boiling the slides in citrate buffer (10 mM sodium citrate, 0.05% Tween 20, pH 6.0) for 20 minutes, followed by gradual cooling to room temperature. Slides were then washed three times with PBS, permeabilized with 0.2% Tween 20 for 20 minutes, and washed again three times with PBS. Blocking was performed using Image-iT™ FX Signal Enhancer (Invitrogen) for 15 minutes. Primary antibody staining was carried out overnight at 4°C using the following antibodies diluted in PBS: rabbit anti-HIF1A (1:100, Invitrogen, PA1-16601), goat anti-NeuN (1:200, Invitrogen, PA5-143586), and chicken anti-GFAP (1:200, EnCor Bio, CPCA-GFAP). For each tissue block, a no-primary antibody control was included to assess non-specific staining. The next day, slides were washed three times with PBS (5 minutes each) and incubated for 1 hour at room temperature with secondary antibodies diluted in PBS: donkey anti-goat AlexaFluor 594 (1:500, Invitrogen, A32758), goat anti-rabbit AlexaFluor 488 (1:500, Abcam, ab150077), and goat anti-chicken IgY AlexaFluor 594 (1:500, Invitrogen, A-11042). When multiplexing donkey anti-goat and goat anti-rabbit antibodies, donkey anti-goat was applied first, followed by three PBS washes, and then goat anti-rabbit was applied to avoid cross-reactivity. Following secondary antibody incubation, slides were washed three times with PBS, stained with DAPI (1:1000), and mounted using Fluoromount-G™ Mounting Medium (Invitrogen). Slides were allowed to cure overnight, and images were acquired using a Leica Stellaris 8 DIVE confocal microscope.

### Statistical Analysis

Unless otherwise specified, the data are presented as the mean ± s.e.m. Student’s two-tailed t-tests and Mann– Whitney U tests were used to assess the significance. P < 0.05 was considered significant. Statistical analyses were performed using Graphpad Prism or R programming language.

## RESULTS

### Compression induces neuronal loss *in vitro*

Solid stress is known to impair neurological function *in vivo*; however, its specific molecular effects on neurons are poorly understood. To investigate this, we used an *in vitro* compression system to apply controlled mechanical compression to human neurons co-cultured with murine astrocytes for 2 months (**Figure 1A**). Using our established protocol (28), we generated induced excitatory neurons (iNs) by forced expression of the transcription factor neurogenin 2 (Ngn2) in human iPSCs from three different sex and ethnic origin donors (referred to as iN #1-3) (**Figure 1B**). We applied 0.14 kPa of compressive stress, mimicking forces exerted by a growing GBM tumor as measured in our prior *in vivo* studies (4). After 24 hours of compression, the iN cultures were stained with propidium iodide (PI) and Hoechst. Live imaging (**Figure 1C**) revealed a more than 2-fold increase in the number and frequency of PI+ cells (indicating loss of membrane integrity consistent with late apoptosis or necrosis) in compressed iNs compared to controls (**Figure 1D-E**), indicating extensive cell death. To specifically assess neuronal viability, we used GFP-expressing iNs to distinguish them from glia. Compressed iN cultures from iPSCs #1 and #2 showed significant neuronal loss (**Figure 1F**) and increased numbers of PI+ neurons (**Figure 1G**). In contrast, GFP- astrocytes did not show signs of increased apoptosis (**Figure S1A**), but total cell numbers increased (**Figure S1B**), suggesting glial proliferation. These results demonstrate that pathophysiological levels of mechanical compression directly lead to neuronal cell death. Among the three iPSC lines, iN #1 neurons consistently exhibited the most robust and reproducible response to compression and thus were selected for further morphological analyses.

### Solid stress reduces the number of synapses and neuronal network activity

Given the pronounced neuronal loss observed under compressive stress, we next investigated whether mechanical compression also compromises synaptic integrity and neuronal network activity. Labeling for synapsin, a presynaptic vesicle marker and MAP2, a marker for postsynaptic dendrites, we analyzed synaptic morphological changes based on three different metrics: synaptic puncta density, puncta area, and puncta intensity, which reflect synapse number and clustering (**Figure 2A-D**). Upon compression, we detected a decrease of all three parameters, indicating a reduction of the number and clustering of synaptic connections. To probe for the functional consequences following mechanical compression, we performed GCaMP8m-based calcium imaging (**Figure 2E-F**). In compressed iNs, we detected a robust decrease in the single-neuron event frequency and amplitude (**Figure 2G-H**) and a less-pronounced decrease in the synchronous event frequency and amplitude (**Figure 2I-J, Movie S1**). This result is indicative of reduced neuronal network activity caused by neuronal and synaptic loss.

**Figure 2.**
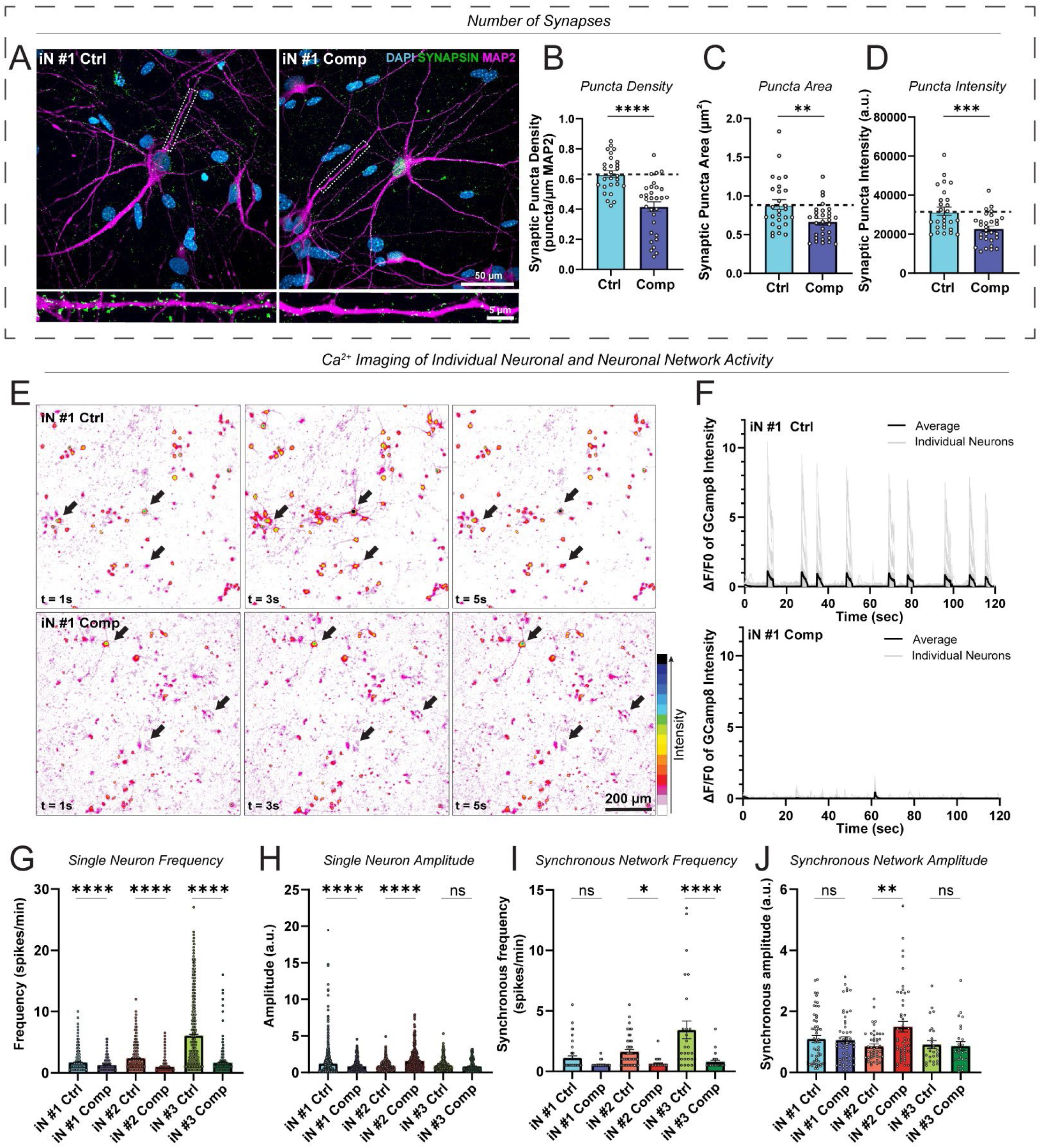
Compression decreases the number of synapses in human neurons. (**A**) Representative confocal images of iN #1 compressed and uncompressed human neurons, showing decreased levels of synaptic puncta. (blue = DAPI, green= Synapsin, magenta = MAP2) (Scale bar = 50μm). (**B-D**) Quantification of synaptic density (p = <0.0001), area (p = 0.0036), and intensity (p = 0.0005) in iN #1. Student’s unpaired t-test n = 27 uncompressed images, n = 30 compressed images, from 2 biological replicates for each condition. (**E**) Representative heat maps for calcium imaging at 2 seconds intervals (Scale bar = 200 μm). (**F**) Representative traces of GCamp8 transients showing fluorescence intensity changes (ΔF/F0) over time. Compressed neurons show decreased event frequency. (**G-J**) Quantification of calcium imaging of iN #1-3 compressed and uncompressed human neurons. Frequency (iN #1 p = <0.0001. iN #2 p = <0.0001. iN #3 p = <0.0001) and amplitude plots (iN #1 p = <0.0001. iN #2 p = <0.0001. iN #3 p = 0.2411) show the activity of individual neurons, while synchronous frequency (iN #1 p = 0.3312, iN #2 p = 0.0121, iN #3 p = <0.0001) and amplitude (iN #1 p = 0.9998, iN #2 p = 0.0032. iN #3 p = >0.9999) show the network activity. One way ANOVA, n = 4 biological replicates in iN #3, n=7 biological replicates in iN #2 and iN #1 per group. Error bars, mean ± SEM.

### Solid stress activates hypoxia response, pro-inflammatory, and apoptosis pathways in human neurons and murine glia

To investigate mechanisms by which solid stress contributes to neuronal dysfunction, we performed RNA-seq on iN/mGlia co-cultures after 24h of mechanical compression. To distinguish between neuronal and glial transcriptional responses, sequencing reads were aligned separately to the human and mouse reference genomes using BBmap (**Figures 3A-B, S1C**). In human iNs, gene set enrichment analysis identified the hypoxia response as the most strongly activated pathway (**Figure 3C**). Leading edge analysis highlighted increased expression of genes involved in glycolysis (*HK2, PDK1, PGK1, ENO1, SLC2A1*), angiogenesis (*VEGFA, ANG, ANGPTL4, CXCL12, CXCR4*), protein synthesis (*BNIP3L, BNIP3, FAM162A*), and apoptosis (*FAM162A, BNIP3, BNIP3L, BBC3, CASP8, BAX*), indicating a mix of pro-survival and pro-apoptotic hypoxia-driven responses (**Figure 3D**). Additionally, we observed significant activation of the neuronal apoptotic program that included both overlapping and distinct markers from the hypoxia response, such as *DAXX, FOXO3, DDIT3 (*CHOP*), PML, ATM, CHEK2, E2F1,* and *PARP3* (**Figure 3D**). Notably, we detected activation of inflammatory signaling pathways, including the NF-κB, AP-1, and JAK/STAT, as evidenced by elevated expression of *RELA, NFKBIA, TNFAIP3, TLR4, CEBPB, MAP3K8, FOS, JUN, SOCS3, IL6ST, IL10RA,* and *OSMR (***Figure 3D**). These findings suggest that neuronal responses to mechanical stress may be compounded by inflammatory signals released from neighboring glia.

**Figure 3:**
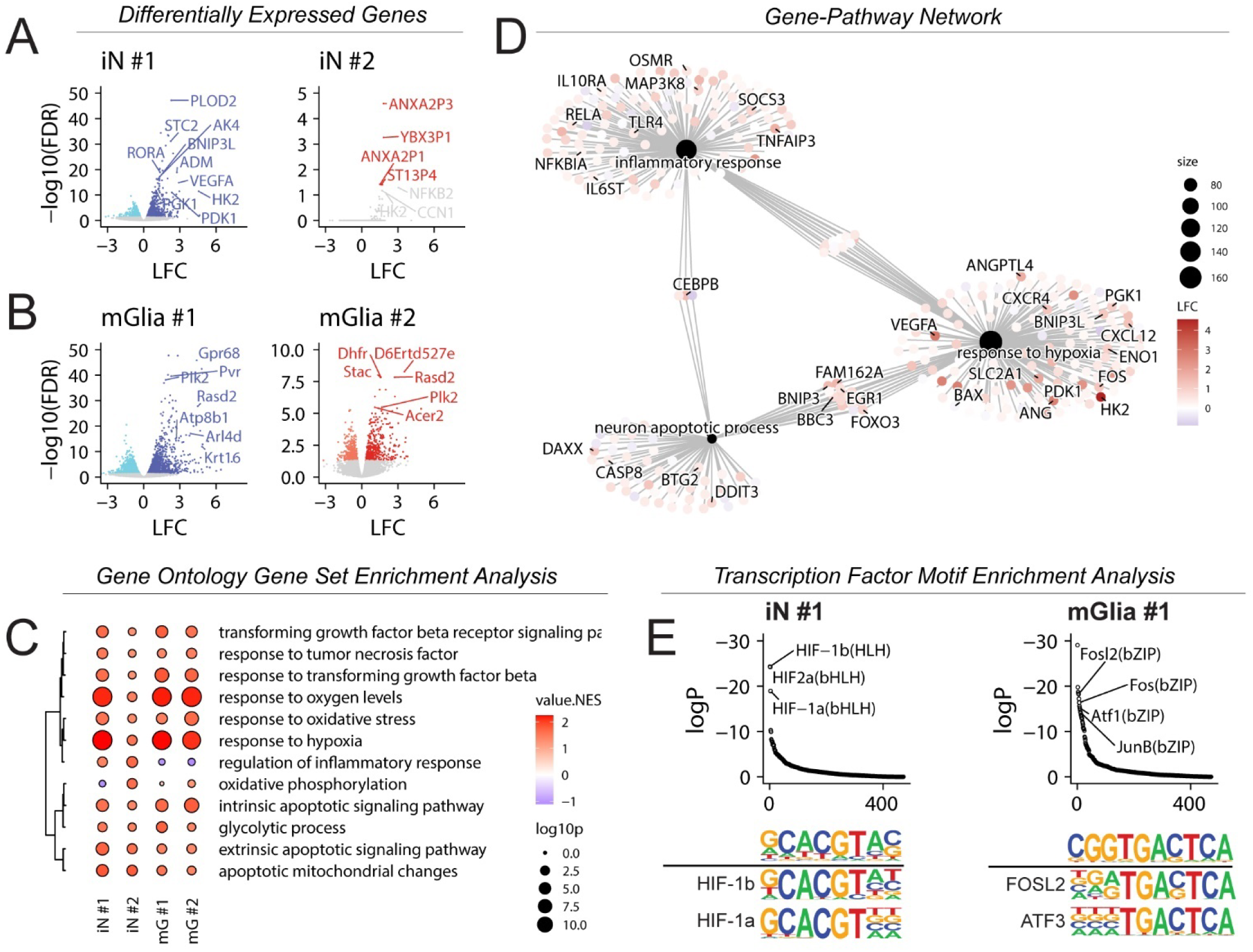
Compression activates hypoxia, apoptosis, and inflammatory pathways in human neurons. Volcano plots for human neurons (iNs) (**A**) and murine glia (mGlia) (**B**) showing up and downregulated genes in dark and light hues, respectively. Top differentially expressed genes are labeled and are enriched in hypoxia response genes; LFC: log fold change. (**C**) Dot plot showing normalized enrichment scores (NES) for iNs and mGlia and select gene ontology (GO) terms. Clustering is based on semantic similarity of GO terms. NES: normalized enrichment score; log10P: negative base 10 logarithm of the adjusted p value. (**D**) GO term to gene graph showing relative expression of leading edge genes for iN #1 in three select pathways. size: number of genes in the leading edge. (**E**) Transcription factor motif enrichment analysis of upregulated (FDR<0.05, LFC>0) genes in iN #1 and mGlia. Motifs are ranked by logP (logarithm of adjusted p value) based on cumulative binomial test. At the bottom, sequence logos for enriched motifs are aligned against known transcription factor motifs for comparison. N=4 biological replicates from 4 experiments for iNs #1 and #2.

In murine glia, mechanical compression similarly activated hypoxia and intrinsic apoptotic pathways (**Figure 3C**). In addition, murine glia exhibited strong upregulation of NF-κB pathway transcription factors (*Fosl1, Fosl2, Jun, Junb, Fosb, Fos, Rel*) and pro-inflammatory cytokines (*Cxcl1, Cxcl2, Cxcl3, Il23a*), consistent with astrocytic and/or microglial activation (astro-/microgliosis) (**Dataset S1**). To further define the glial cell types contributing to this response, we performed cell type deconvolution using the Tabula Muris single-cell RNA-seq atlas (41). This analysis identified astrocytes as the top-scoring cell type (**Figure S1D**).

To identify putative upstream regulators driving these responses, we conducted transcription factor motif enrichment analysis. In neurons, HIF-1 and HIF-2 binding motifs were strongly enriched in transcription start sites of upregulated genes, consistent with elevated hypoxia signaling (**Figure 3E**). In glia, AP-1 transcription factor motifs were significantly enriched, suggesting that glial inflammatory response may be driven in part by AP-1 signaling. Together, these data suggest that mechanical compression activates HIF-1, inflammatory, and apoptotic signaling in both neurons and glia. Glia-derived inflammation may contribute to neuronal injury, potentially compounding the direct effects of solid stress.

### Solid stress activates reactive glial gene expression programs in human astrocytes and microglia

Astrocytes can become reactive in response to both direct mechanical injury and signals released by stressed or dying neurons (42). To determine whether solid stress can directly trigger astrogliosis in the absence of neurons, we subjected immortalized human astrocytes (IHAs) to *in vitro* compression. After 24 hours, compressed astrocytes showed increased expression of GFAP––a canonical marker of reactive astrocytes (**Figure 4A**).

**Figure 4:**
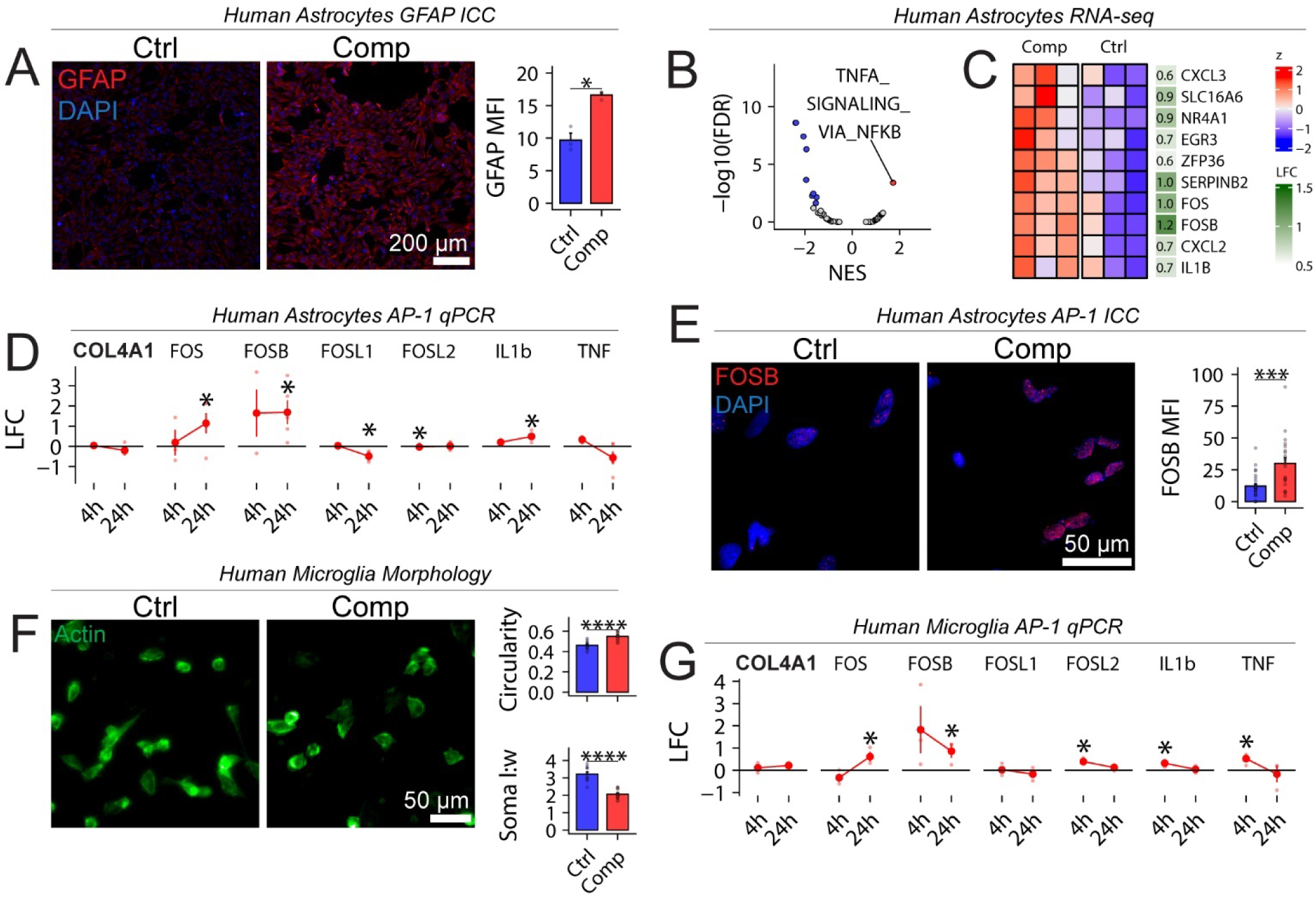
Mechanical compression activates astro- and micro-gliosis via upregulation of the AP-1 signaling pathway. (**A**) Immunofluorescence analysis of GFAP, a marker of reactive astrocytes, in compressed immortalized human astrocytes. (n = 3 biological replicates per group from one experiment). (**B**) Volcano plot showing top dysregulated pathways in astrocytes based on gene set enrichment analysis of MSigDB HALLMARK gene sets; FDR: false discovery rate; NES: normalized enrichment score. (**C**) Normalized and scaled expression of top differentially expressed genes in astrocytes from the HALLMARK_TNFA_SIGNALING_VIA_NFKB gene set. LFC: log fold change. (**D**) qPCR gene expression quantification of NF-κB and AP-1 pathway genes in compressed astrocytes; n = 3 biological replicates for the 4h time point and n = 5 biological replicates for the 24h time point. (**E**) Immunofluorescence analysis of FOSB in compressed astrocytes (n = 4 biological replicates per group from one experiment). (**F**) Changes in morphology of C20 immortalized human microglia are quantified as circularity and soma length:width ratio using ImageJ based on actin staining (n = 3 biological replicates per group from one experiment). (**G**) qPCR gene expression quantification of NF-κB and AP-1 pathway genes in compressed microglia (3 biological replicates for all timepoints). Error bars, mean ± SEM. All p values are based on Student’s t-tests. Stars indicate significance levels: * p < 0.05, ** p < 0.01, *** p < 0.001, **** p < 0.0001. For (**D, G)** p values are calculated by comparing comp. v.s. ctrl at each timepoint. Log fold change (LFC) is calculated using ddCt method; dCt is calculated relative to GAPDH expression.

To explore the mechanisms driving this response, we performed RNA-seq on compressed IHAs. Gene set enrichment analysis (GSEA) revealed significant activation of the NF-κB signaling pathway (**Figure 4B-C**), including immediate-early AP-1 transcription factors *JUN*, *FOS*, and *FOSB*, similar to the response observed in murine glia. To assess the dynamics of this response, we performed qPCR at 4 and 24 hours post-compression. *FOS* and *FOSB* levels increased rapidly at 4 h and remained elevated at 24 hours (**Figure 4D**), indicating sustained activation rather than a transient, acute response. This was further confirmed by immunocytochemistry, which showed increased nuclear translocation of Fos-B under compression (**Figure 4E**). IL1B was specifically induced at 24 hours, further supporting a chronic inflammatory response. Notably, TNF-α––a key upstream activator of canonical NF-κB signaling––was not upregulated, pointing instead to a non-canonical, potentially mechanosensitive mode of NF-κB activation.

*In vivo*, astrogliosis may also result from indirect effects of solid stress, such as hypoxia resulting from vascular compression, as we have shown previously (4). To determine whether changes in local oxygen tension contributed to astrocyte activation, we repeated the compression experiment under hypoxic conditions (1% O2) and found that compression still induced upregulation of GFAP (**Figures S2A-B**). To further rule out the effects of residual hypoxia that may persist even in 1% O2, we used the HypoxiTRAK™ oxygen-sensing probe, which showed no increase in biologically relevant hypoxia levels under compression (**Figures S2C-D**). Consistently, HIF1A expression remained unchanged (**Figures S2E-F**), supporting the conclusion that compression-induced astrogliosis occurs independently of hypoxia.

Because astrocyte reactivity often occurs in concert with microglial activation, we next asked whether microglia also respond directly to compressive stress. Microglia are highly sensitive to tissue damage and can initiate neuroinflammatory cascades that influence both astrocyte reactivity and neuronal survival (43). To assess their response, we examined the effects of compression on the morphology of immortalized human C20 microglia (32–34), which changes rapidly in response to injury (44). Morphological analysis revealed that compressed C20 microglia adopted a more reactive, amoeboid phenotype, with increased circularity and reduced soma length-to-width ratio (**Figure 4F**) (44). Similar to astrocytes, compressed microglia showed sustained upregulation of FOS and FOSB, but their expression of pro-inflammatory cytokines TNF-α and IL1B was transient, returning to baseline by 24 hours (**Figure 4G**). These results indicate that mechanical compression directly induces reactive transcriptional programs in both astrocytes and microglia via NF-κB signaling. This glial activation may amplify neuroinflammation and contribute to neuronal injury and death under solid stress.

### Solid stress upregulates hypoxic and inflammatory gene programs *in vivo* in murine and human brains

Taken together, the above results demonstrate that mechanical compression directly induces neuronal dysfunction by activating pro-apoptotic and hypoxia response pathways in neurons and inducing reactive phenotypes in astrocytes and microglia. To assess the relevance of these findings *in vivo*, we applied chronic, localized mechanical compression to the mouse brain using a custom-designed apparatus (**Figure 5A**). In compressed cortical regions, we observed loss of NeuN+ neurons that spatially coincided with the formation of a dense GFAP+ astroglial scar (**Figure 5B,D,E**), suggesting astrocyte-mediated neuronal toxicity. Multiplex immunostaining revealed increased nuclear HIF1A expression in surviving neurons within compressed regions (**Figure 5C,F,G**), indicating activation of hypoxia signaling in response to mechanical stress.

**Figure 5.**
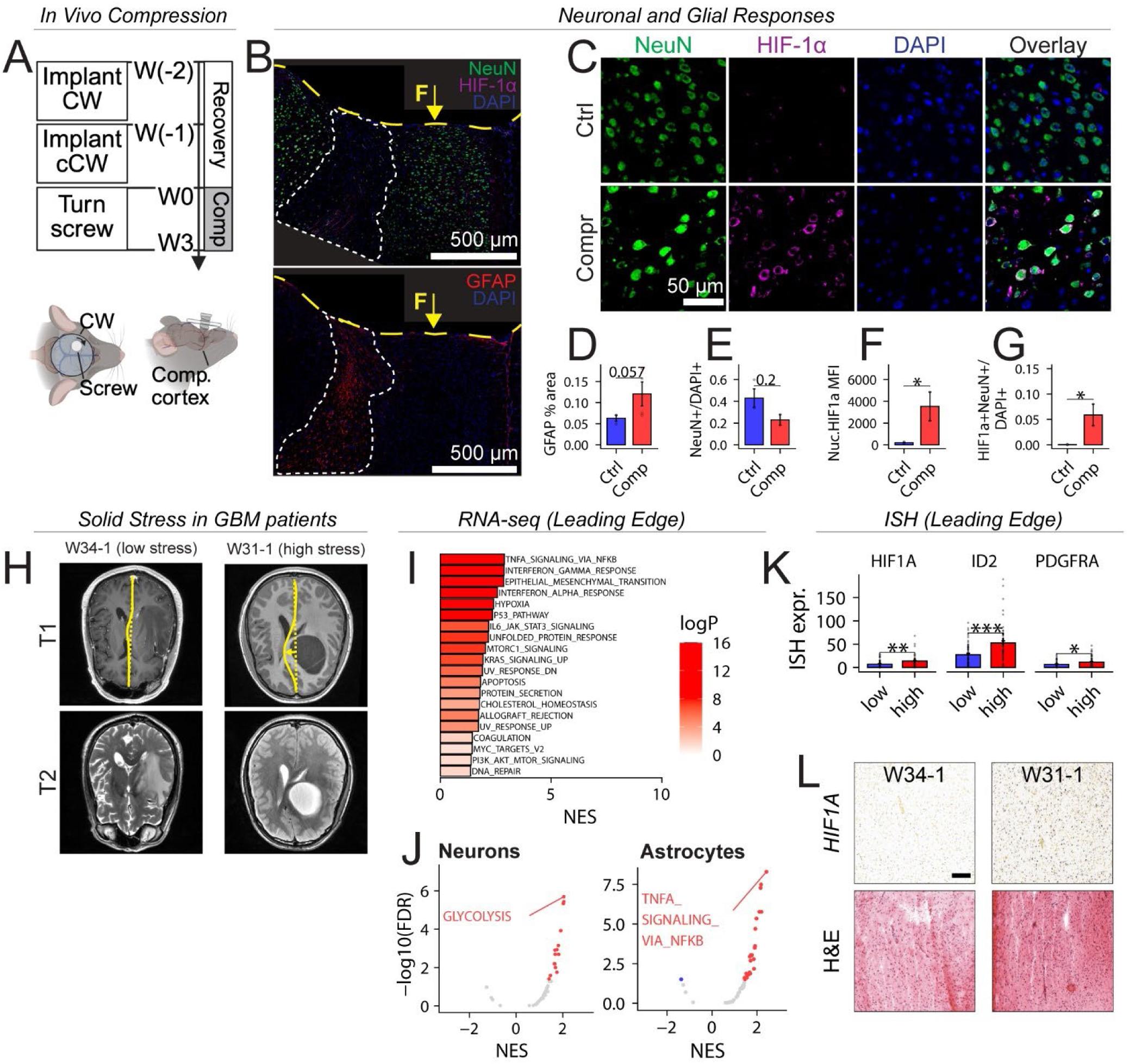
Solid stress promotes hypoxic and inflammatory gene expression programs *in vivo*. (**A**) Schematic and timeline of the *in vivo* cortical compression system. The compression device consists of a set screw mounted on a cranial window, separated from the brain surface by a biocompatible, deformable membrane. The screw gradually expands the compression volume at approximately 1.3 mm³/day over 3 weeks, mimicking the growth rate of nodular intracranial GBM models (4). (**B**) Representative immunofluorescence images of cortices subjected to mechanical compression, stained for HIF-1α and NeuN (top) or GFAP (bottom). A region lacking NeuN signal, indicative of neuronal loss, and showing increased GFAP staining, consistent with reactive astrogliosis, is outlined with a dashed white line. The area of cortical indentation (compression divot) is marked with a yellow dashed line, and the site of applied compression is indicated by a yellow arrow; F: compressive force. (**C**) Immunofluorescence images showing NeuN and HIF-1α expression in the compressed cortex (Scale bar: 50 µm). The bottom row (Comp) shows higher-magnification insets taken from the compressed region highlighted in panel B. (**D-G**) Quantification of immunohistochemistry data shown in panels B and C. GFAP % area is quantified as fraction of pixels above an expression intensity threshold; neuron frequency (NeuN+/DAPI+) is quantified as the number of NeuN+ particles divided by the number of DAPI+ particles. Nuclear HIF1a mean fluorescence intensity is quantified for Neun+DAPI+ nuclei. The frequency of HIF1a- expressing neurons (HIF1a+NeuN+/DAPI+) is quantified as the number of HIF1a+NeuN+ particles divided by the number of DAPI+ particles. Mann–Whitney *U* test using n = 2 biological replicates for Comp and n = 3 biological replicates for Ctrl from 1 experiment. (**H**) Representative MRI images from GBM patients with high (W31-1) and low (W34-1) estimated solid stress, showing T2-weighted (bottom) and T1 post-contrast (top) scans. (**I**)Top dysregulated pathways identified by gene set enrichment analysis (GSEA) of IvyGAP RNA-seq data from histologically normal “leading edge” regions from patients with high estimated solid stress. NES: normalized enrichment score. logP: negative logarithm of adjusted P value. (**J**) GSEA of BayesPrism- deconvolved bulk RNA-seq data reveals activation of glycolysis and NF-κB signaling pathways in neurons and astrocytes in regions of high solid stress. (**K**) Quantification of in situ hybridization (ISH) data in the leading edge region. “Low” and “High” refer to patients with low and high estimated solid stress, respectively. Student’s two- tailed t-test using n = 9 biological replicates for “high” and n = 10 for “low” groups. (**L**) Representative HIF1A ISH and H&E micrographs showing increased HIF1A expression in the histologically normal regions in the patient with high estimated solid stress (also shown in panel H).

To determine whether similar patterns occur in human disease, we analyzed data from the Ivy Glioblastoma Atlas Project (45). For each patient, we quantified midline shift (MLS) on T1 post-contrast MRI (**Figure 5H**), a widely used radiographic marker of mass effect that correlates with poor neurological outcomes in GBM (46, 47) as well as stroke (48–51) and TBI (52–56). Because MLS reflects both solid stress and peritumoral edema, we regressed MLS on edema volume and used the residuals to stratify patients by estimated solid stress (see **Methods, Figure S3**).

Differential expression analysis of RNA-seq data from the peritumor of these patients revealed upregulation of NFκB, p53, interferon signaling, and hypoxia response pathways in high-stress patients (**Figure 5I**). *In situ* hybridization (ISH) data confirmed elevated *HIF1A* expression in the same regions, along with increased levels of oligodendroglial precursor markers *ID2* and *PDGFRA* (**Figure 5J**). To further investigate cell-type-specific responses at the tumor margin, we performed BayesPrism deconvolution of IvyGap bulk RNA-seq data using a published scRNA-seq reference dataset (38). Gene set enrichment analysis revealed increased glycolysis in neurons and elevated NF-κB signaling in astrocytes, specifically in patients with high solid stress (**Figure 5K- L**)—closely mirroring our experimental findings. Together, these data support a model in which tumor-induced solid stress activates HIF1A-mediated stress responses in neurons and NF-κB–driven inflammation in astrocytes. This combination of hypoxic-like stress and glial reactivity may contribute to peritumoral neurotoxicity and neurological impairment in GBM patients.

## DISCUSSION

We previously demonstrated that chronic compression of the mouse cortex—at a rate similar to GBM growth— leads to neuronal loss and neurological deficits (4). However, the molecular mechanisms linking compression to these outcomes remained unclear. In this study, we used multiscale compression models to mimic the mechanical forces exerted by GBM and examine their direct impact on neurons and glial cells. Our findings demonstrate that compression leads to a marked decrease in the frequency of spontaneous calcium transients in neurons, indicating disrupted intrinsic network activity. This loss of network synchrony, along with decreased synaptic density, may underlie the cognitive impairments, motor deficits, and elevated seizure risk observed in GBM patients (57, 58).

Mechanistically, compression appears to reduce iN network activity by driving apoptosis and decreasing the number and size of functional synapses (59). We specifically probe for the number of synapses and size of synaptic clusters per neuron, though further work is needed to determine whether synaptic dysfunction precedes neuronal loss or vice versa. In some neurological diseases, decreased synaptic activity itself can contribute to neuronal loss (59). For instance, in Alzheimer’s Disease, the onset of synaptic dysfunction occurs prior to cortical neuron death, suggesting that a reduction in synaptic activity can precipitate cell death (60).

On the molecular level, we detect robust activation of the HIF-1 signaling pathway in compressed neurons, with known pro- and anti-apoptotic effects in the context of neurologic diseases (61, 62). In some models, HIF-1α stabilization has been shown to promote recovery in models of spinal cord injury (63), cerebral ischemia (64–66), and traumatic brain injury (67). However, other studies suggest that HIF-1α can directly contribute to secondary injury (62, 68). The hypoxia response signature observed in our data includes genes involved in anaerobic metabolism and angiogenesis, consistent with an adaptive role for HIF-1α. However, we also observe upregulation of pro-apoptotic genes, including *BNIP3* and *BNIP3L* (NIX)––both regulated by HIF-1α and known to induce mitophagy and cell death (69, 70). This aligns with our previous observation of increased LC3-positive vesicles in compressed mouse brains (4), suggesting that compression may trigger neuronal death through HIF- 1α/BNIP3-mediated mitophagy. Given the pleiotropic functions of HIF-1α, our bulk RNA-seq data cannot resolve whether individual neurons activate both pro- and anti-apoptotic pathways simultaneously, or if distinct subpopulations activate these programs independently. Future studies incorporating single-cell RNA-seq may help address this question.

HIF-1 signaling pathway is classically associated with cellular responses to hypoxia, although oxygen-independent, including mechanosensitive modes of HIF-1 activation have been demonstrated previously (71). *In vivo*, the direct effects of solid stress are confounded by reduced blood perfusion, making it difficult to disentangle oxygen-dependent from oxygen-independent mechanisms of HIF-1 activation. Our *in vitro* compression device overcomes this challenge through the use of oxygen-equilibrated agarose hydrogels and the transwell system, which allow for gas and nutrient exchange under compression. Using this system, we find that HIF-1 response pathways are induced by mechanical compression even in the absence of overt hypoxia, pointing to a mechanosensitive mode of HIF-1 activation.

In addition to its direct effects on neurons, we observe that compression activates the pro-inflammatory and mechanosensitive (72) AP-1 pathway in both murine mixed glial cultures and human astrocyte and microglia cell lines. Compressed glia upregulate cytokines such as *Cxcl1*, *Cxcl2*, *Cxcl3*, *Il23a*, *IL1B*, and *TNFA* which are known to contribute to neuroinflammation and may exacerbate neuronal injury (73–75). Notably, while both astrocytes and microglia activate the immediate-early AP-1 transcription factors *FOS* and *FOSB*, they exhibit distinct temporal dynamics. Microglia respond rapidly—within hours—to compression by transiently upregulating transcripts for TNF-α and IL-1β, cytokines with known neurotoxic roles (43); whereas astrocyte reactivity appears to be more sustained. We also observe species-specific responses: murine glia show upregulation of hypoxia response pathway, whereas human astrocytes did not upregulate hypoxia-related genes or show nuclear translocation of HIF1A. This discrepancy may reflect the use of immortalized human cell lines, which do not fully recapitulate the behavior of primary glia. Future work using iPSC-derived astrocytes might better model the physiological responses of human glia and help clarify whether the observed differences are due to species-specific biology.

Our mechanistic insights are further supported by *in vivo* and patient data. In *in vivo* compression models, we observe elevated neuronal HIF1A activity and astrocytic inflammation in cortical areas exposed to compression. Interestingly, we observe these effects in regions adjacent to, rather than directly beneath, the site of applied pressure. This suggests that both compressive and tensile forces—present in these adjacent areas—contribute to the observed cellular responses. In GBM patients, we estimate solid stress using midline shift (MLS)––a well-recognized radiographic marker of mass effect. Because MLS is strongly influenced by increased fluid pressure due to peritumoral edema, we regress MLS on edema volume and use the residuals as a surrogate for solid stress. However, this approach relies on the assumption that the part of the midline shift (MLS) not accounted for by edema directly represents solid stress, which may overlook other biomechanical and anatomical factors that contribute to the observed mass effect. In future studies, solid stress should be directly measured using established techniques (76, 77) and combined with unbiased transcriptomic profiling of peritumoral regions to more accurately dissect the effects of solid stress on the surrounding neural tissue in patients.

In this study, we use Ngn-2-induced human neurons - purely excitatory neurons, as they produce robust action potentials and reliably form functional synapses (28). Nevertheless, we observe significant variability in how neurons derived from different iPSC lines respond to compression. For example, neurons obtained from neonatal iPSCs #3 do not exhibit a significant increase in apoptosis observed in the iPSCs #2 and iPSCs #1 neurons. This variability may stem from intrinsic genetic or epigenetic differences between donor iPSC lines. In TBI, for example, genome-wide association studies have shown that common genetic variations substantially contribute to inter-patient variability in clinical outcomes (78). While our sample size is limited, these observations underscore the need for further investigation into the role of genetic variation in neuronal response to injury.

In summary, our findings indicate that chronic compression triggers a cascade of direct and indirect signaling events that ultimately lead to neuronal death, damage, and disruption of neuronal network activity. These findings point to potential therapeutic opportunities. For instance, our previous work has shown that lithium can partially rescue neuron loss and restore function under similar conditions (4). Our iN-based *in vitro* compression model provides a useful platform for screening neuroprotective compounds. The mechanistic insights from this study offer a foundation for developing interventions that target neuronal dysfunction in GBM and may also be applicable to other conditions involving mechanical brain injury, such as TBI or tumefactive MS.

## ACKNOWLEDGEMENTS

This work was supported by the National Institutes of Health (NIH/NIGMS R35GM151041, awarded to M.D.) and the Harper Cancer Research Institute through the Cancer Cure Venture Grant (awarded to M.D. and C.P.). Additional support was provided by the Berthiaume Institute for Precision Health Graduate Fellowship and the Scientific Artificial Intelligence Fellowship (both awarded to M.Z.). RNA-seq library preparation and sequencing was performed by the Genomics & Bioinformatics Core Facility at the University of Notre Dame. We gratefully acknowledge Prof. Felipe H. Santiago-Tirado for providing the C20 microglia cells. High-performance computing resources were provided in part by the Notre Dame Center for Research Computing. Histological analyses were conducted at the Notre Dame Histology Core, and fluorescence imaging was performed at the Notre Dame Integrated Imaging Facility.

## Contributions

Conceptualization: M.Z., A.W., M.D., C.P.

Data curation: M.Z., A.W., J.N., J.L., J.M.

Formal analysis: M.Z., A.W., J.M.,

Funding acquisition: M.Z., M.D., C.P.

Investigation: M.Z, A.W., J.L., J.M., J.B-P., B.B., M.P., R.R., A.B.

Methodology: M.Z., A.W., J.M., C.P.

Project administration: C.P., M.D.

Resources: M.D., C.P.

Software: M.Z., A.W., C.M. Supervision: M.D.. C.P.

Validation: M.Z., A.W., C.P.

Visualization: M.Z., A.W., C.P.

Writing - original draft: M.Z., A.W., M.D., C.P.

Writing - review & editing: M.Z., A.W., J.N., J.L., J.M., J.B-P., C.M., B.B., M.P., R.R., A.B., H.S., B.S., M.D., C.P.

