## Supplementary material for "Mechanical compression induces neuronal apoptosis, reduces synaptic activity, and promotes glial neuroinflammation in mice and humans": Suppl Figures

**Supplementary figures and figure captions**

**Movie S1.** Calcium live imaging of control iN #1 and compressed iN #1. Both compressed and control iNs were grown in culture for 60 days. On Day 59, the compressed iNs were compressed for 24 hours and then imaged with the control iNs. iNs were imaged in an imaging buffer for 2 minutes with a 50 msec framerate. Scale bar = 100 µm.

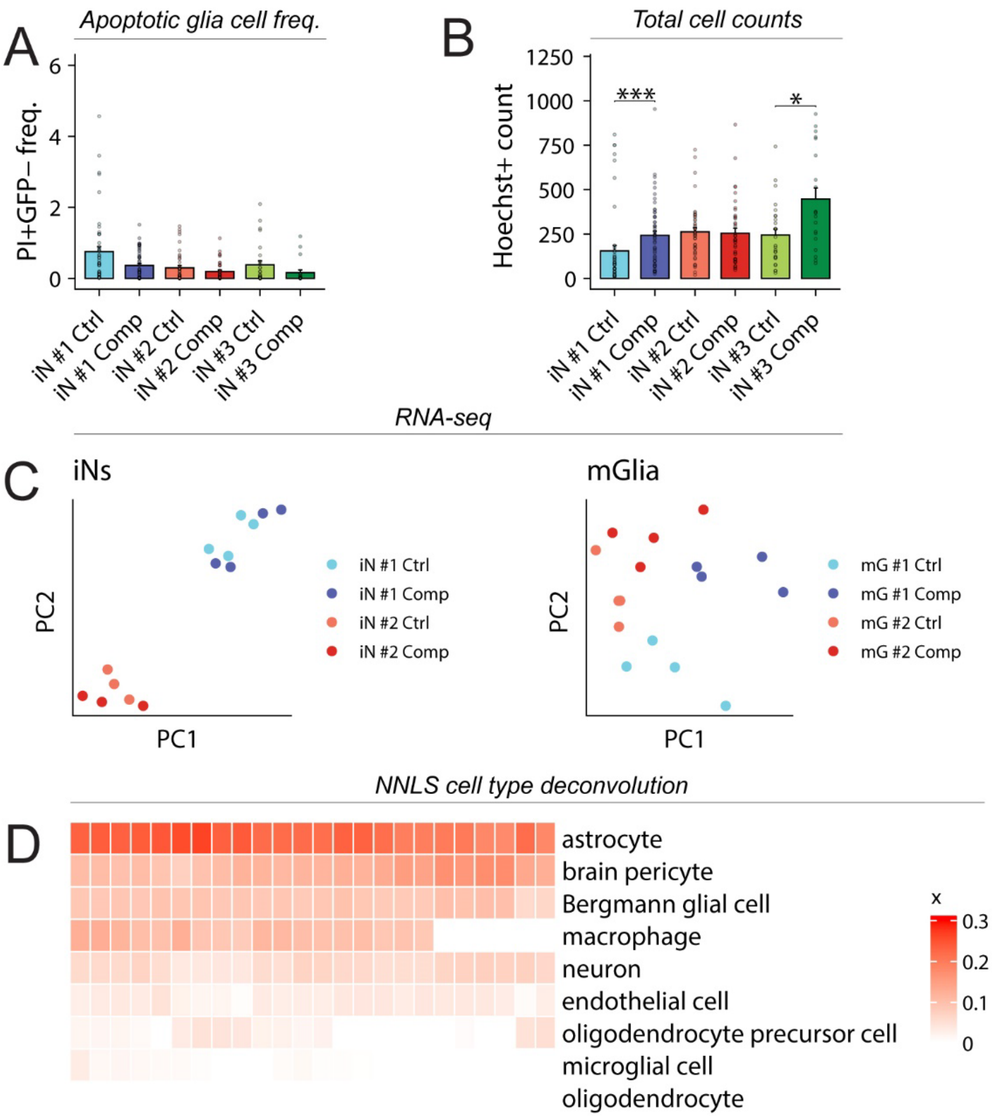


**Figure S1. Quantification of glial cell viability and molecular characterization of neuronal and glial populations.** (**A**) Apoptotic glial cell frequency is quantified by counting PI+GFP- particles per field of view (FOV) and dividing the count by the number of Hoechst+ particles (**B**). Total cell numbers are quantified as the number of Hoechst+ particles per FOV. (**C**) Principal component analysis of human neurons and murine glia shows that iN transcriptomes separate based on source iPSC donor, while murine glia separate based on experimental condition. (**D**) Non-negative least squares (NNLS)-based cell type deconvolution of bulk RNA-seq data from murine glial cultures reveals strong enrichment of the astrocyte gene expression signature. Mann–Whitney U test with Holm–Bonferroni adjustment for multiple comparisons using N = 4 biological replicates for iNs #3, n=7 biological replicates in iNs #2 and iNs #1 neurons per group; the data are based on 7 independent experiments; Error bars: mean ± SEM. Due to puncta-based quantification, some frequencies may exceed 1. Statistical significance is indicated as follows: * = P ≤ 0.05;  *** = P ≤ 0.001.

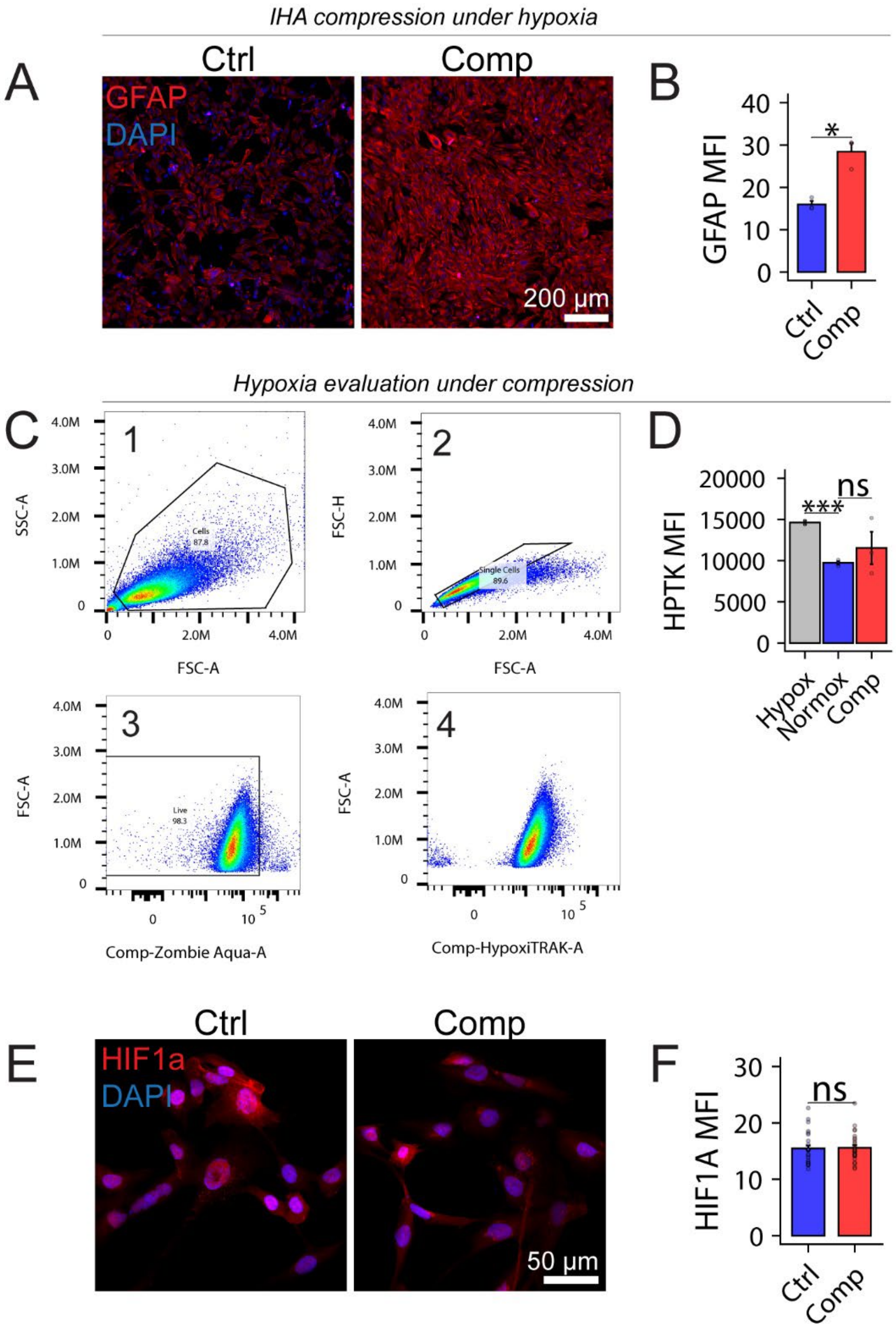


**Figure S2. Astrocyte activation in response to compression occurs independently of hypoxia.** (**A–B**) Immunofluorescence staining of GFAP in immortalized human astrocytes (IHA) cultured under hypoxia (1% $O_2$), with or without compression (n = 3 biological replicates per group). (**C–D**) Flow cytometry gating strategy and quantification of HypoxiTRAK fluorescence in human astrocytes cultured under normoxia (20% $O_2$), hypoxia (1% $O_2$), or compression. Compression does not induce biologically significant hypoxia. (**E–F**) Immunofluorescence analysis of HIF1A shows no increase in nuclear HIF1A accumulation in compressed astrocytes.

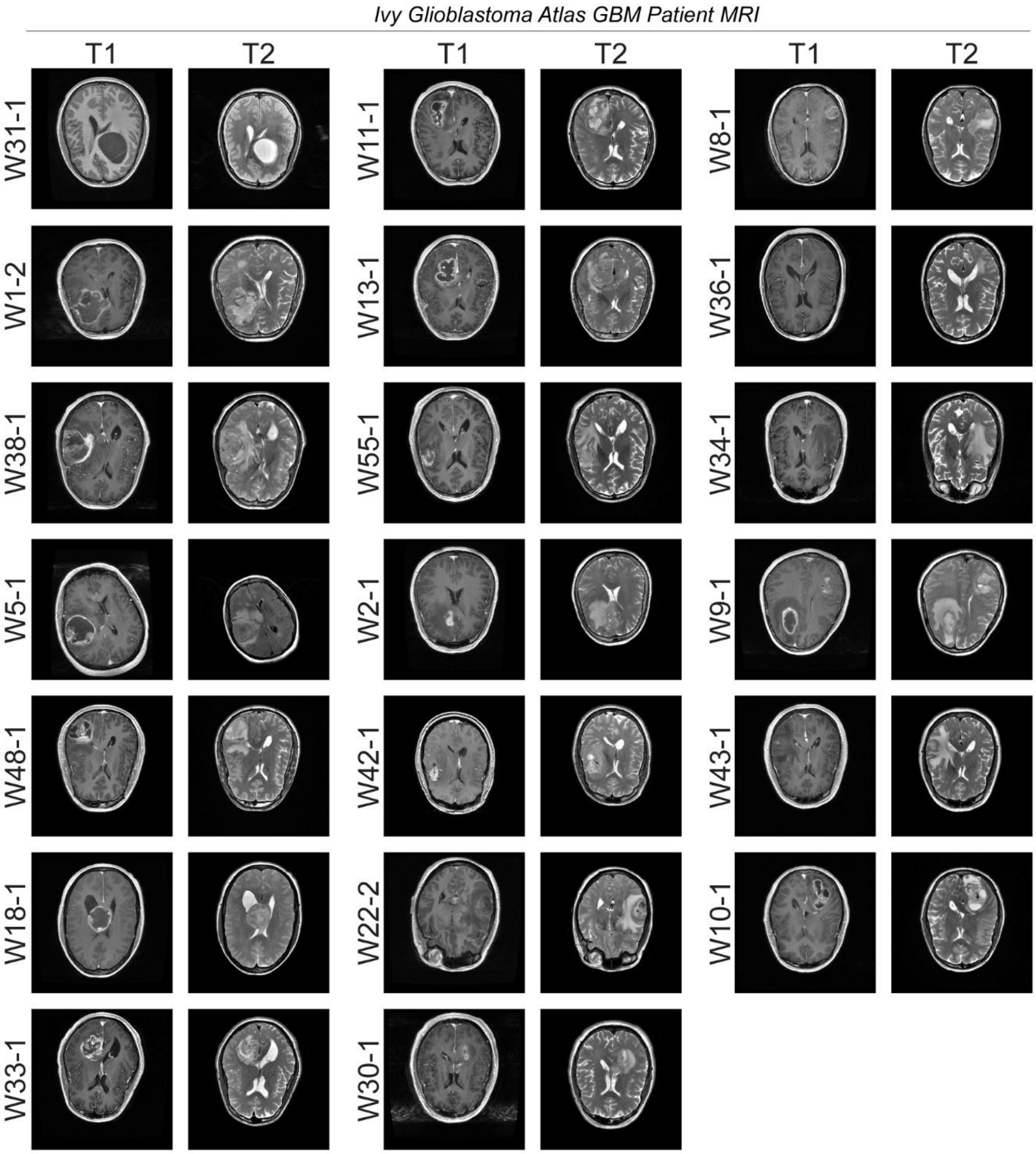


**Figure S3. Solid stress estimation in the Ivy Glioblastoma patient cohort.** Patients are ranked by estimated solid stress levels (top-to-bottom, then left-to-right). T1-weighted and T2-weighted MRI images are shown for each patient to illustrate the contributions of  tumor burden and peritumoral edema.
